# Generation of a novel SARS-CoV-2 sub-genomic RNA due to the R203K/G204R variant in nucleocapsid: homologous recombination has potential to change SARS-CoV-2 at both protein and RNA level

**DOI:** 10.1101/2020.04.10.029454

**Authors:** Shay Leary, Silvana Gaudieri, Matthew D. Parker, Abha Chopra, Ian James, Suman Pakala, Eric Alves, Mina John, Benjamin B. Lindsey, Alexander J Keeley, Sarah L. Rowland-Jones, Maurice S. Swanson, David A. Ostrov, Jodi L. Bubenik, Suman Das, John Sidney, Alessandro Sette, COVID-19 Genomics UK (COG-UK) consortium, Thushan I. de Silva, Elizabeth Phillips, Simon Mallal

**Author notes:** Corresponding author Prof. Simon Mallal. These authors contributed equally to this work (Shay Leary, Silvana Gaudieri, Matthew D. Parker). These authors also contributed equally to this work (Thushan I. de Silva, Elizabeth Phillips, Simon Mallal).

## Abstract

**Background:** Genetic variations across the SARS-CoV-2 genome may influence transmissibility of the virus and the host’s anti-viral immune response, in turn affecting the frequency of variants over-time. In this study, we examined the adjacent amino acid polymorphisms in the nucleocapsid (R203K/G204R) of SARS-CoV-2 that arose on the background of the spike D614G change and describe how strains harboring these changes became dominant circulating strains globally.

**Methods:** Deep sequencing data of SARS-CoV-2 from public databases and from clinical samples were analyzed to identify and map genetic variants and sub-genomic RNA transcripts across the genome.

**Results:** Sequence analysis suggests that the three adjacent nucleotide changes that result in the K203/R204 variant have arisen by homologous recombination from the core sequence (CS) of the leader transcription-regulating sequence (TRS) rather than by stepwise mutation. The resulting sequence changes generate a novel sub-genomic RNA transcript for the C-terminal dimerization domain of nucleocapsid. Deep sequencing data from 981 clinical samples confirmed the presence of the novel TRS-CS-dimerization domain RNA in individuals with the K203/R204 variant. Quantification of sub-genomic RNA indicates that viruses with the K203/R204 variant may also have increased expression of sub-genomic RNA from other open reading frames.

**Conclusions:** The finding that homologous recombination from the TRS may have occurred since the introduction of SARS-CoV-2 in humans resulting in both coding changes and novel sub-genomic RNA transcripts suggests this as a mechanism for diversification and adaptation within its new host.

## Introduction

It is believed SARS-CoV-2 originated from a bat coronavirus transmitted to humans, likely via an intermediate host such as a pangolin, acquiring a furin-cleavage site in the process. This new motif allows cleavage at the boundary of the S1 and S2 domains of the spike protein in virus-producing cells (1). A SARS-CoV-2 variant in the spike protein, D614G (B.1 lineage), emerged early in the epidemic and has rapidly became dominant in virtually all areas of the world where it has circulated (2). Several studies have shown this variant to be associated with higher viral RNA levels in the upper respiratory tract, higher titers in pseudoviruses in-vitro (2, 3) and increased infectivity (4, 5). More recently, emerging lineages from this genetic background (B.1.1.7 – ‘Alpha or UK variant’, B.1.351 – ‘Beta or South African variant’, or B.1.617.2 - ‘Delta variant’) have been identified with reported rapid local expansions of these viruses.

The diversification of coronaviruses can occur via point mutations and recombination events (6, 7) that can result in increased prevalence due to selective advantage related to increased infectiousness and transmission of the virus or by chance. Evidence of viral adaptation to selective pressures as a virus spreads among diverse human populations has important implications for the ongoing potential for changes in viral fitness over time, which in turn may impact transmissibility, disease pathogenesis and immunogenicity. Furthermore, the functional impact of new genetic changes need to be considered in the performance of diagnostic tests, ongoing public health measures to contain infection around the world and the development of universal vaccines and antiviral therapies including monoclonal antibodies.

Here we examined a variant of SARS-CoV-2 that emerged within the subset of sequences harboring the D614G variant and contains three adjacent nucleotide changes spanning two residues of the nucleocapsid protein (R203K/G204R; B.1.1 lineage) that has resulted in a novel sub-genomic RNA transcript. Sequence analysis suggests these changes are the result of homologous recombination from the core sequence (CS) of the leader transcription-regulating sequence (TRS). This event introduced a new TRS between the RNA binding and dimerization domains of nucleocapsid providing the template for the generation of a novel sub-genomic RNA transcript. Further novel sub-genomic RNA transcripts arising in association with incorporation of leader sequence and TRS were also observed, suggesting homologous recombination from this region as a potential mechanism for SARS-CoV-2 diversification and adaptation within its new host.

## Methods

### Study Design

This study utilized deposited SARS-CoV-2 genomic sequences in public databases, with a further 981 Oxford Nanopore Technology genomes and clinical metadata from Sheffield, UK, as a validation set, to identify and map genetic variants and sub-genomic RNA transcripts across the genome. Accession numbers and links to datasets are in Supplementary Material.

### SARS-CoV-2 sequence generation from patients with COVID-19

SARS-CoV-2 sequences, with matched clinical metadata, were generated using samples taken for routine clinical diagnostic use from 981 individuals presenting with COVID-19 disease to Sheffield Teaching Hospitals NHS Foundation Trust, Sheffield, UK. This work was performed under approval by the Public Health England Research Ethics and Governance Group for the COVID-19 Genomics UK consortium (R&D NR0195). Following extraction, samples were processed using the ARTIC Network SARS-CoV-2 protocol. After RT-PCR, SARS-CoV-2 specific PCR and library preparation with Oxford Nanopore LSK-109 and barcoding expansion packs NBD-104 and NBD-114 samples were sequenced on an Oxford Nanopore GridION X5 using R9.4.1D flow cells. Bases were called with either fast or high accuracy guppy with demultiplexing enabled and set to --require-both-ends. Samples were then analyzed using ARTIC Network pipeline v1.1.0rc1.

### SARS-CoV-2 sequence acquisition from public repositories

Complete SARS-CoV-2 genome sequences were downloaded from the GISAID EpiCoV repository on 24^th^ January 2021 (https://www.gisaid.org/). The complete dataset of 455,774 sequences with coverage across the genome were aligned in CLCbio Genomics Workbench 12 (QIAGEN Bioinformatics) to the GenBank reference sequence NC_045512.2. Aligned sequences were exported in FASTA format and imported into Visual Genomics Analysis Studio (VGAS), an in-house program for visualizing and analyzing sequencing data (http://www.iiid.com.au/software/vgas). The chronological appearance of the sequences was generated using the collection dates for each of the sequences. Of note, our current knowledge of the global circulating variants is dependent on the ability of laboratories in different countries to deposit full genome length SARS-CoV-2 sequences and may be subject to ascertainment bias. As such, the frequencies of specific variants shown may not reflect the size of the outbreak. However, the data does provide the opportunity to predict the presence of specific variants in areas given the known epidemiology within different countries and regions. A subset of subjects also had individual deep sequence reads deposited in the Sequence Read Archive (SRA) at www.ncbi.nlm.nih/sra. These sequence reads were downloaded and aligned as indicated above.

### Identification of amino acid substitutions

Codon usage output allowed for identification of amino acid substitutions across the SARS-Cov-2 genome. A cut-off of 5% frequency within the consensus SARS-CoV-2 protein sequences was set to obtain the codon usage across all sequences and as shown in S1 Table. The viral polymorphisms detected are present in viral variants sequenced using different NGS platforms (e.g. nanopore, Illumina) and the Sanger-based sequencing method making it unlikely that the new changes are sequence or alignment errors. In addition, different laboratories around the world have deposited sequences with these polymorphisms in the database and examination of individual sequences in the region failed to uncover obvious insertions/deletions likely representing alignment issues or homopolymer slippage.

### HLA peptide binding prediction

The region containing the adjacent amino acid polymorphisms in the nucleocapsid was divided into sliding windows of 8-14 amino acids. NetMHC 4.0 (http://www.cbs.dtu.dk/services/NetMHC/) and NetMHCpan 4.0 (http://www.cbs.dtu.dk/services/NetMHCpan/) with default settings were utilized to predict HLA-class I binding scores and binding differences across all HLA class-I alleles for the original 2003 SARS and current SARS-CoV-2 sequences harboring the R203/G204 and K203/R204 polymorphisms in the nucleocapsid (output listed in S2 Table).

### HLA peptide binding assays

MHC was purified from the Steinlin EBV transformed homozygous cell line (IHWG ID: 9087; A*01:01, B*08:01 and C*07:01) using the B123.2 (anti-HLA-B, C) and W6/32 (anti-class I) monoclonal antibodies, and classical MHC-peptide inhibition of binding assays performed, as previously described (8). To develop an HLA C*07:01-specific binding assay, the IEDB was utilized to identify candidate peptides reported as HLA-C*07:01 epitopes or eluted ligands. One peptide (3424.0028; sequence IRSSYIRVL, Macaca mulatta and Homo sapiens DNA replication licensing factor MCM5 289-297) was radiolabeled and found in direct binding assays to yield a strong signal with as little as 0.5 nM MHC. Subsequent inhibition of binding assays established that 3424.0028 bound with an affinity of 0.21 nM. To establish that the putative assay was specific for C*07:01, and not co-purified B*08:01, two additional peptides previously reported as HLA-C*07:01 ligands were also tested, with one found to bind with high affinity (IC50 67 nM) and the other with intermediate (IC50 1600 nM). At the same time, a panel of known B*08:01 ligands were not found to have the capacity to inhibit binding of radiolabeled 3424.0028 (S3 Table). By contrast, when the same panel of peptides was tested in the previously validated B*08:01 assay (9), 3424.0028 was found to bind with about 1500-fold lower affinity, all of the known B*08:01 ligands bound with IC50s <10 nM, and the C*07:01 ligands with affinities >1000 nM.

### Sub-genomic RNA classification & quantification in the Validation Dataset

We developed a tool, “periscope” (v0.0.0), to classify and quantify sub-genomic RNA in the Sheffield ARTIC network Nanopore dataset (10). The tool can be downloaded from git-hub at https://github.com/sheffield-bioinformatics-core/periscope. Briefly, this tool uses local alignment to identify putative sub-genomic RNA supporting reads and uses genomic reads from the same amplicon to normalize.

### RNA structure modeling

The RNAfold program from the ViennaRNA Web Server (http://rna.tbi.univie.ac.at/) was used for structural predictions using the default settings and the minimum free energy structures were acquired using the base-pairing probability color scheme. The Dot-bracket folding notations were obtained for each of the R203K/G204R sequences and used for Junction Explorer (nature.njit.edu/biosoft/Junction-Explorer/) and CHS-align (nature.njit.edu/biosoft/CHSalign/).

### Statistical Analysis

Fisher exact test was used to compare the proportion of subjects with specific sub-genomic RNA transcripts. P values less than 0.05 was used as the statistical threshold. Comparisons between sub-genomic and genomic RNA expression in R203/G204 compared to K203/R204 containing sequences was made using the Mann-Whitney U test, corrected for multiple comparisons using the Holm method. Logistic and linear regression modeling used to explore the impact of K203/R204 and other co-variates on hospitalization, CT values and sub-genomic RNA expression.

## Results and Discussion

### Adjacent nucleocapsid polymorphisms emerged from the existing spike protein D614G variant

We utilized publicly available SARS-CoV-2 sequences from the GISAID database (available on the 24^th^ of January 2021; www.gisaid.org) to identify amino acid polymorphisms arising in global circulating forms of the virus in relation to region and time of collection. Of the 455,774 circulating variants there were 29 amino acid polymorphisms present in >5% of the deposited sequences (of a total of 9413 sites; S1 Table) including the spike D614G variant (B.1 lineage) that emerged early in the pandemic and the adjacent R203K/G204R variants (B.1.1 lineage) in the nucleocapsid protein (11) that formed one of the main variants emerging from Europe in early 2020. As of the end of January 2021, the K203/R204 variant comprises 37.4% of globally reported SARS-CoV-2 sequences (Fig 1) and almost exclusively occurs on the D614G genetic background (S4 Table).

**Fig 1.**
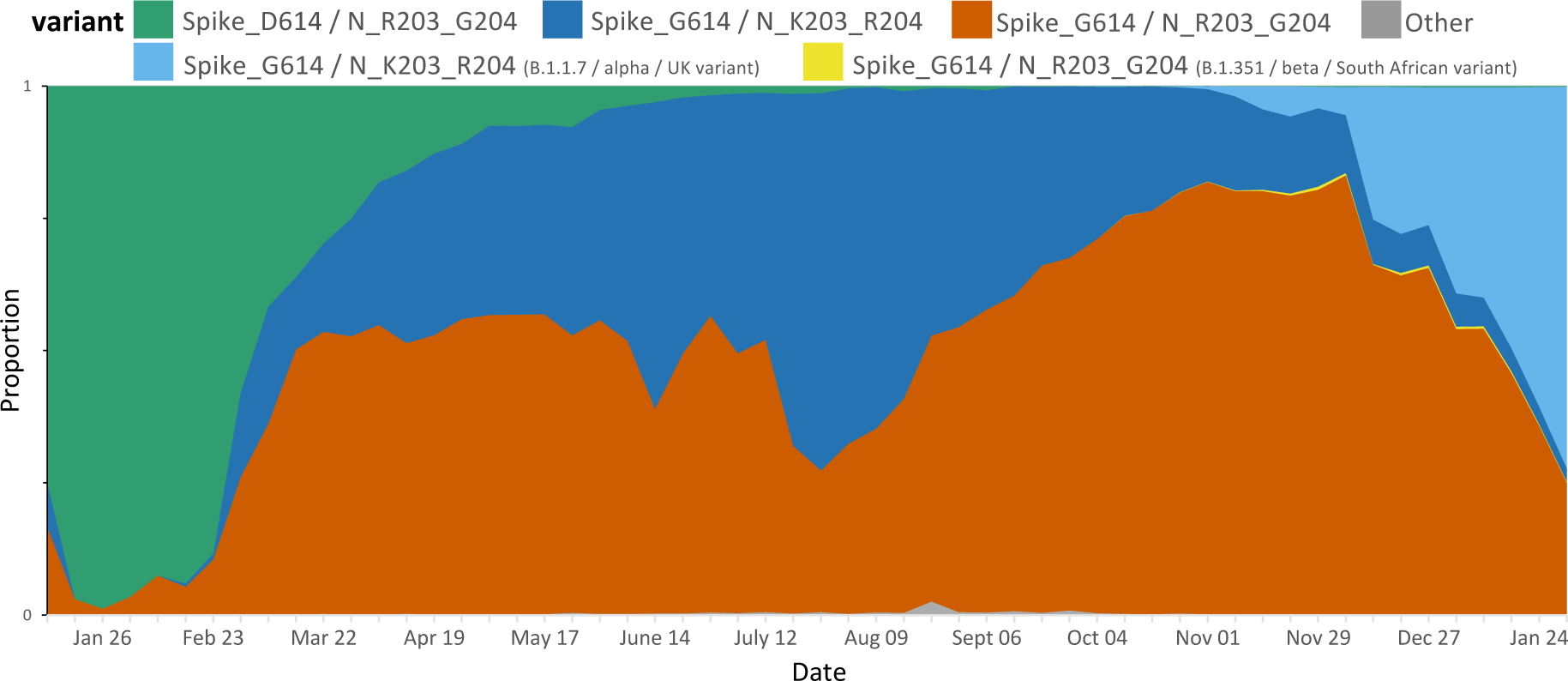
Proportion of weekly deposited SARS-CoV-2 sequences globally (n=455774). The D614G (B.1) variant has become one of the dominant forms globally. Note a small proportion of deposited sequences did not include information regarding specific collection date and as such were excluded.

Although the D614G change rapidly increased in prevalence in almost all regions, the prevalence rates of the K203/R204 subset of the D614G variant are variable in different geographic areas and over-time (Fig 2). For example, an almost complete replacement of D614 by G614 was noted in South America between March and April 2020 and a similar replacement pattern was seen with the K203/R204 variant most marked in Chile, Argentina and Brazil (12). A closer examination of the deposited sequences in the UK shows the K203/R204 variant increasing in prevalence early in 2020 but the second wave later in the year shows a shift in the proportion of deposited sequences with the R203/G204 subset of the D614G variant (B.1.177 lineage) until the recent appearance of the B.1.1.7 ‘Alpha or UK variant’ that harbors the K203/R204 polymorphisms (S1 Fig and S4 Table); supporting a likely increased infectivity of this variant.

**Fig 2.**
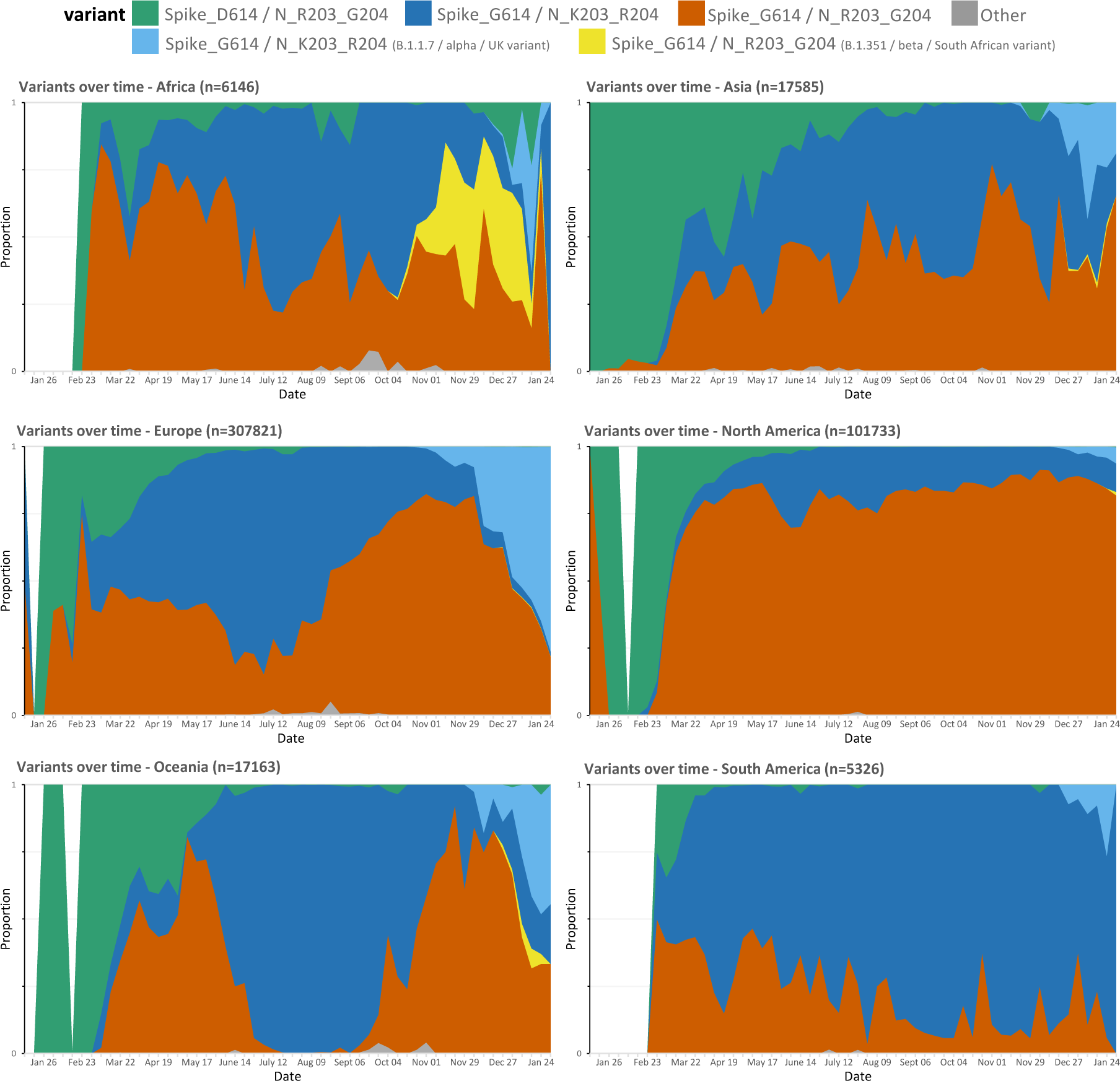
Proportion of weekly deposited SARS-CoV-2 sequences by region. The proportion of R203/G204 to K203/R204 sub-variants of the D614G variant differs in different regions with recent increases in the frequency of new variants.

### Amino acid polymorphisms due to three adjacent nucleotide changes in the nucleocapsid likely due to homologous recombination

Of the publicly available sequences examined with the two amino acid polymorphisms K203/R204, all showed the three adjacent nucleotide changes from A<u>GG G</u>GA to A<u>AA C</u>GA. There was no differential codon usage for the K203/R204 variant in the database. However, there was evidence of low frequency alternative codon usage for arginine at 203 (AGA) for the R203/G204 variant and for lysine (AAG) at 203 for the K203/G204 variant (S5 Table). Overall, circulating variants that contain the intermediate codon as the consensus that could facilitate a single step from the AGG arginine codon to the AAA lysine codon at position 203 appear rare among captured variants to date (S5 Table). Furthermore, a K203 polymorphism alone was seen in 0.3% and an R204 polymorphism alone seen in only 0.02% of sequences (S5 Table). The low frequency K203/L204 and K203/P204 variants are both one nucleotide step from the K203/R204 variant, have been deposited into the public databases (November 2020) well after the emergence of the K203/R204 variant (February 2020) and accordingly likely arose from this genetic background.

The rapid emergence of closely linked polymorphisms in viruses can also reflect strong selection pressure on this region of the genome in which the original mutation incurred a replicative capacity, or other fitness cost, which could be restored by a linked compensatory mutation. Evidence for such adaptations with closely linked compensatory mutations are known to occur under host immune pressure as is well established for other RNA viruses such as HIV (13–15) and Hepatitis C virus (16). In the absence of anti-viral treatment, these viruses have such a high rate of viral replication, error-prone polymerases and lack associated proofreading, mismatch repair, and other nucleic acid repair pathways generating a swarm of viral variants with ongoing recombination between variants (in the case of HIV) being generated continuously. As a result, selection pressure exerted by immune responses or other selective pressures effectively operate on each separate residue independently (15). In contrast, coronaviruses encode proofreading machinery and have a propensity to adapt by homologous recombination between viruses (6) rather than necessarily by classic stepwise individual mutations driven by selective pressures effectively operating on individual viral residues. Furthermore, a simulation based on the nucleocapsid genomic region and allowing up to 10 random mutations indicates the likelihood of observing three consecutive nucleotide changes is less than 0.0005. These findings argue against stepwise change of the nucleotides for the R203K/G204R variant.

The introduction of the AAACGA motif by homologous or heterologous recombination is a more parsimonious mechanistic explanation and would have immediately resulted in both an R to K change and adjacent G to R change at the positions 203 and 204, respectively. It is critical to determine if the introduction of the AAACGA motif has induced any replicative or other fitness change for the virus as a result of either structural or functional changes in the RNA or the concomitant change of amino acids from R203/G204 to K203/R204 and any related structural or functional impact on the nucleocapsid protein.

### SARS-CoV-2 itself as likely source for homologous recombination

To identify possible viral sources for homologous recombination with SARS-CoV-2, we initially performed a search of the motif in the nucleocapsid in related beta coronaviruses from human and other species in the public databases and only found the presence of the R203/G204 combination. We performed a similar search in our metatranscriptome data generated from a cohort study consisting of 65 subjects of whom 43 had acute respiratory infections and 22 were asymptomatic. From the data we assembled near complete and coding complete viral genomes of the Coronavirus (NL63 - alpha, OC43 - beta, 229E - alpha), RSV (A, B), Rhinovirus (A, B, C), Influenza (A - H3N2), and Bocavirus family. None of the alpha coronaviruses had the R203/G204 or K203/R204 combination or indeed any variation at these sites (n=14; sequence depth >3000). We then performed a search for stretches of similarity using varying window sizes (>14 base-pair (bp) including the motif) in all sequences. A 14bp window was selected as 14bp has been shown to be the minimum amount of homology required for homologous recombination in mammalian cells (17). No significant hits were identified. However, the AAACGA sequence encoding the K203/R204 amino acids overlaps with the CTAA<u>ACGAAC</u> motif of the leader transcription-regulating sequences (TRS; core underlined) (18) of SARS-CoV-2 itself and this core sequence motif is also found near the start codon of the protein for surface glycoprotein (S), ORF3a, E, M, ORF6, ORF7a, ORF8, ORF10 and nucleocapsid, in keeping with its known roles in mediating template switching and discontinuous transcription (18).

### Deep sequencing confirms quasi-species with the leader sequence linked to known or introduced TRS region

Discontinuous transcription of SARS-CoV-2 results in sub-genomic RNA (sgRNA) transcripts containing 5’-leader sequence-TRS-start codon-ORF-3’. These RNA transcripts should also be captured from reads generated from NGS platforms. We therefore reasoned we should be able to find such sequences within deep sequencing reads at the sites of known sub-genomic regions (corresponding to the ORFs) and adjacent to position 203/204 of the nucleocapsid in subjects infected with the K203/R204 variant but not in those with the R203/G204 variant (Fig 3).

**Fig 3.**
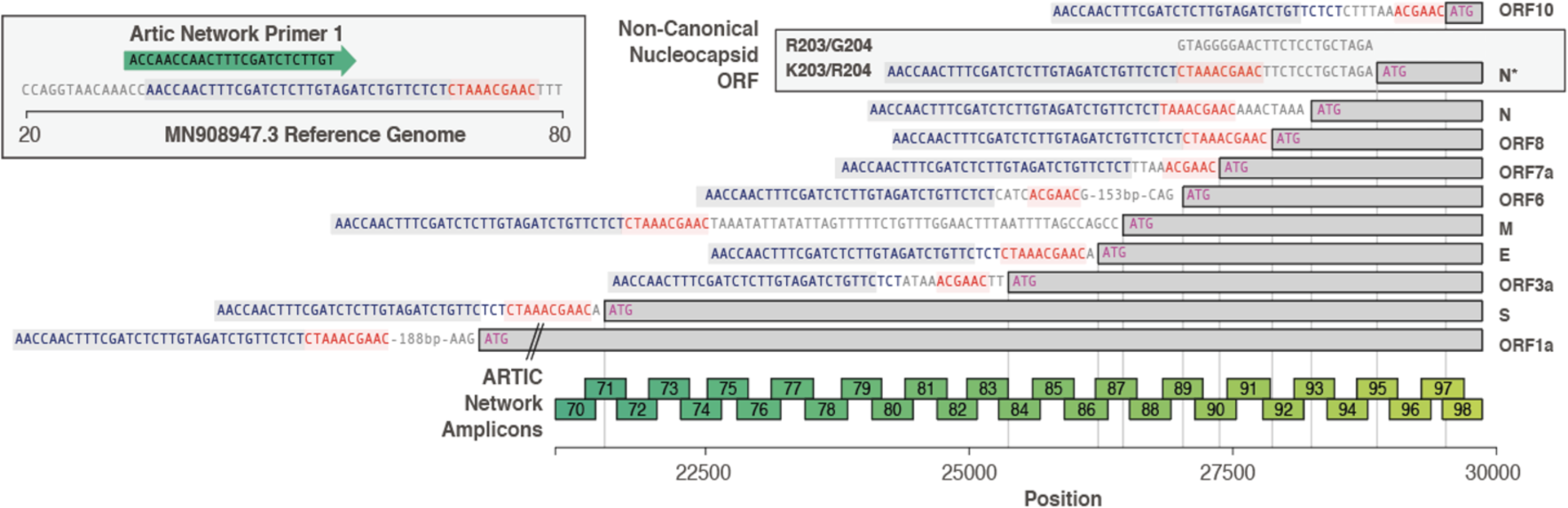
The configuration of canonical sgRNAs and the novel non-canonical nucleocapsid sgRNA (N*) in SARS-CoV-2. The bottom bar illustrates the presence of the leader sequence (blue text) followed by the transcription-regulating sequence (TRS; red text) within the genomic sequence that continues into the first ORF 1a. The presence of other canonical sgRNA transcripts in which the leader sequence and TRS precede the start codon (methionine; pink) of the other proteins are shown. The presence of the novel non-canonical sgRNA transcript containing the K203/R204 polymorphisms (N*) is shown. The ARTIC primer locations and resultant amplicons are shown.

We searched for sgRNAs in sequence data generated from n=981 patients with COVID-19 based on the ARTIC network protocol (www.artic.network/ncov-2019; Fig 3) and subsequent Nanopore sequencing in Sheffield, UK. As expected, the most frequent sgRNA transcripts in each subject, irrespective of variant, corresponded to the known regions containing the start codon of the SARS-CoV-2 proteins (Fig 4A). However, out of a total of 550 K203/R204 sequences, 231 had evidence (>=1 read containing leader sequence at the novel TRS site) of the non-canonical nucleocapsid sgRNA (42%) but only 1 out of a total of 431 R203/G204 subjects had evidence of the novel sgRNA (likely a false positive as described in S2 Fig).

**Fig 4.**
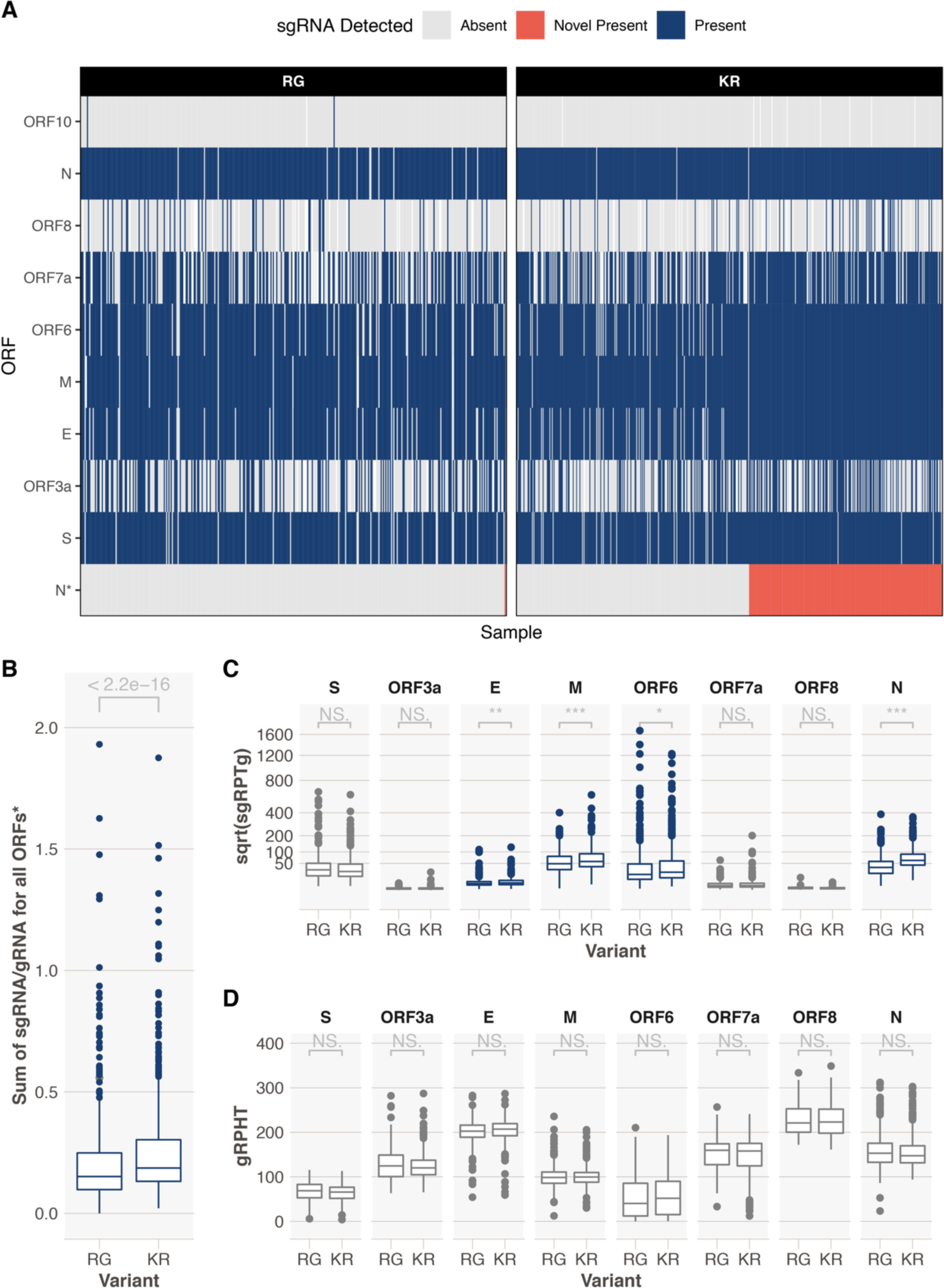
Exploration of sgRNAs in 981 samples from Sheffield, UK. A. A heatmap showing presence or absence of sgRNAs from different ORFs. K203/R204 (KR)-containing sequences have evidence of the novel truncated N ORF sgRNA (N*, red, 233/553, 42%). An ORF sgRNA was deemed present if we could find >=1 read in support. Heatmap is ordered by the presence or absence of the novel sgRNA. There were a total of 448 R203/G204 (RG)-containing sequences and 1 had evidence of a novel sgRNA (likely false positive, Fig S2). **B**. Significantly higher (Mann-Whitney U p < 2.2e-16) total sgRNA in KR-containing compared to RG-containing sequences. **C**. Sub-genomic RNA is significant increased in KR-containing compared to RG-containing sequences for a number of ORFs, most notably nucleocapsid (N; Mann-Whitney U p = 2.06e-37 corrected for multiple testing using the Holm method). Y-axis denotes square root transformed sub-genomic reads normalized to 100,000 genomic reads from the same ARTIC amplicon. **D.** There is no difference in genomic RNA levels (normalized to total mapped reads) between KR- and RG-containing sequences. *novel sgRNA, ORF10 and ORF1a are excluded from this analysis due to ORF10 not being expressed, difficulty in discriminating ORF1a sgRNA from genomic RNA and the novel truncated N sgRNA is only being present in KR-containing sequences. **\*\*\* < 0.001, ** < 0.01, * < 0.05.** All p values shown are following correction for multiple testing with the Holm method.

We confirmed the presence of the novel non-canonical nucleocapsid sgRNA in 27/45 individuals with the K203/R204 variant but in none of 45 individuals with the R203/G204 variant (Fisher test, p=5.0e-11; S6 Table) from the sequence read archive (SRA) database (www.ncbi.nlm.nih/sra). Interestingly, we also found the presence of 23 other non-canonical sgRNA transcripts with the 5’-leader-TRS-start codon-3’ at low frequency in the 90 subjects (irrespective of variant) due to multiple adjacent changes to the consensus sequence across the genome generating new core TRS motifs (including with minor mismatches) (S6 Table). It should be noted that none of these changes are present in the consensus sequence of the SARS-CoV-2 genomes downloaded and represent low frequency quasispecies within individuals. It does, however, suggest other instances of the introduction of the core sequences from the leader TRS elsewhere in the SARS-CoV-2 genome.

### SARS-CoV-2 viruses with K203/R204 are not associated with greater hospitalization with COVID-19 or higher virus levels in the upper respiratory tract

The same dataset from COVID-19 patients in Sheffield, UK, was used to explore whether the K203/R204 variant had any association with clinical outcome. The median age of this cohort was 54 years (IQR 38 to 74) and 59.8% were female. Of these, 440 (44.9%) were hospitalized COVID-19 patients and 42 (4.3%) subsequently required critical care support. A multivariable logistic regression model including 203/204 status, age and sex showed no association of K203/R204 with hospitalization (OR 0.82, 95% confidence intervals (CI 0.58 – 1.16), p=0.259). As expected, higher age and male sex were significantly association with hospitalization with COVID-19 (OR 1.09, 95% CI 1.08 – 1.11, p <2e-16 for age and OR 4.47, 95% CI 3.13 – 6.43, p=2.91e-16 for male sex). Male sex, but not age or 203/204 status, was associated with risk of critical care admission (S7 Table).

We explored whether K203/R204 was associated with greater virus levels in the upper respiratory tract as estimated by cycle threshold (CT) values from the diagnostic RT-PCR. As day of illness will impact CT value, we focused on a subset of the cohort (n=478) where this information was available (all non-hospitalized patients, median symptom day 3, range 1 – 13 days). Data were analyzed with sequences stratified by spike 614 and nucleocapsid 203/204 status (D614/R203/G204, G614/R203/G204 and G614/K203/R204). Multivariable linear regression models showed no impact of G614/K203/R204 compared to G614/R203/G204 status on CT values (p= 0.83, S6B Table), but as expected, later day of symptom onset was significantly associated with higher CT values, therefore lower viral load (S8 Table, p=2.05E-05). Consistent with recent findings (2), presence of a spike D614G variant was significantly associated with lower CT values (higher viral loads) in the same subset of individuals, even when day of illness at sampling is included in the model (S8A Table, D614/R203/G204 vs G614/R203/G204, p=0.00011, Fig 5A & B).

**Fig 5.**
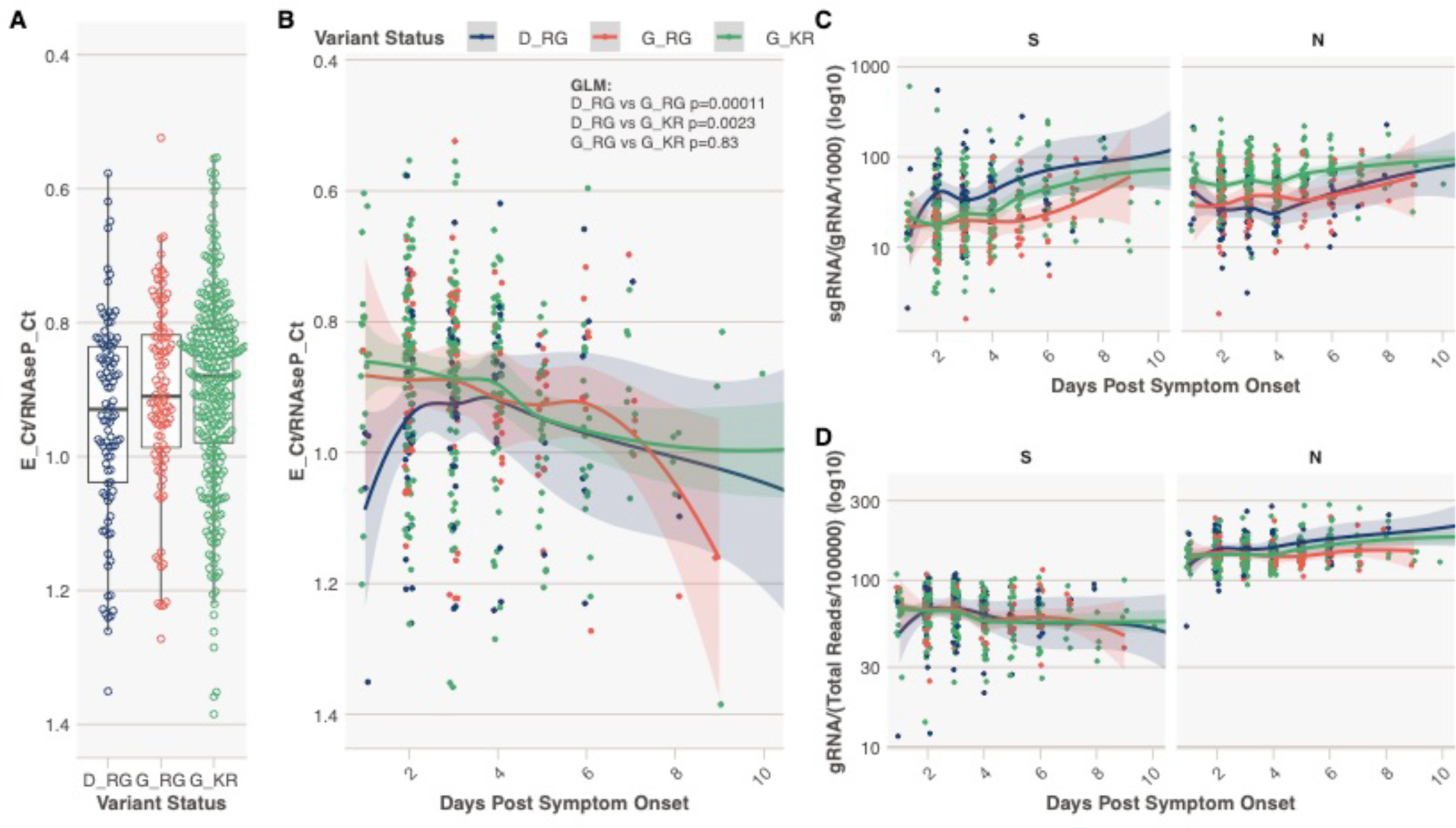
Spike 614 and Nucleocapsid 203/204 Status, Diagnostic Metrics and level of sub-genomic and genomic RNA. A. **A.** E gene cycle threshold (CT) normalized to RNAseP CT stratified by variant status in N = 478 individuals from Sheffield dataset with day of symptom onset data available. This normalization was done to combine and display E gene CT data from two different extraction protocols. Y-axis reversed to aid interpretation, as lower normalized CT values equal higher virus levels. **B.** Normalized E gene CT vs the day of sampling from day of symptom onset. P values provided are from a generalized multivariable linear regression model (GLM) for the difference in normalized E gene CT value between samples containing each variant, with extraction method and day of illness included in the model (Table S6) **C.** Normalized (per 1000 genomic reads) sgRNA levels for ORFs S and N. **D.** Normalized (per 100,000 mapped reads) genomic RNA levels for ORFs S and N.

### SARS-CoV-2 viruses with K203/R204 have evidence of higher sub-genomic RNA expression

We hypothesized that the amount of sgRNA at each of the ORF TRS positions in the SARS-CoV-2 genome in ARTIC nanopore sequencing data could serve as a proxy for expression levels of each of the ORFs due to their positions in the amplicons (Fig 3). To test this hypothesis we developed a tool, periscope (19), which quantifies the number of sgRNA and genomic RNA reads at each ORF TRS position in ARTIC network nanopore sequencing data. We applied periscope to the 981 sequences in the Sheffield validation dataset. To control for the sequencing depth differences evident between amplicons, we determined the amplicon that shares the 3’ primer with the sgRNA reads and used the total count of genomic RNA at this amplicon to calculate the proportion of sgRNA for each ORF. The N ORF sgRNA is expressed at high levels in all samples. ORF10 sgRNA was absent as others have shown (20). A significant increase in sgRNA levels for several ORFs in samples with K203/R204 compared to R203/G204 samples is apparent (Fig 4B). N is the most striking example (Fig 4C, Mann-Whitney U test p value, adjusted for multiple testing p = 2.06e-37), but sgRNA from ORFs E, M and ORF6 are also significantly increased. There is no significant difference in genomic RNA levels (Fig 4D, normalized to total mapped reads) between these two groups.

As discussed above, the K203/R204 variants appear to have emerged within the subset of SARS-CoV-2 sequences with a D614G variant in the spike protein, which has been associated with infections with a higher viral load in the upper respiratory tract. To explore whether the differences between K203/R204 and R203/G204 sequences in sgRNA quantities were due to D614 compared to G614 variant differences, we repeated the comparisons following further stratification of sequences. Interestingly, G614/R203/G204 variants showed *lower* total sgRNA expression than D614/R203/G204 samples (S3 Fig). Of note, sgRNA for spike (S), membrane (M) and envelope (E) ORFs were significantly higher in samples with D614/R203/G204 compared to those with G614/R203/G204 (adjusted p values 1.02e-4 for S, 0.0495 for M and 0.00696 for E). Total sgRNA in G614/K203/R204-containing samples was still significantly higher than in G614/R203/G204 samples (S3A Fig, Mann-Whitney U test p value, adjusted for multiple testing p = 3.5e-6). Similar increases in some individual ORF sgRNA quantities in G614/K203/R204 compared to G614/R203/G204 sequences were also seen, most notably for nucleocapsid (S3B Fig, adjusted p value 1.34e-12).

To ensure that the increase in sgRNA in K203/R204-containing sequences was not due to confounding by differences in sampling date compared to date of symptom onset, we evaluated the impact of K203/R204 and day of illness on sgRNA expression in a multivariable linear regression model using the subset of 478 sequences described above (stratified by D614/R203/G204, G614/R203/G204 and G614/K203/R204 status). Higher sgRNA levels were significantly associated with later day from symptom onset (S9 Table, p=9.9E-08). G614/R203/G204 compared to D614/R203/G204 was again associated with a reduction in sgRNA levels (p=0.011, S9A Table), whereas a K203/R204 change on the background of spike G614-containing sequences was associated with a significant increase in sub-genomic RNA (p=4.51E-05, S9B Table). Spike canonical sub-genomic RNA was higher in D614/R203/G204 samples, whereas nucleocapsid canonical sub-genomic RNA was higher in G614/K203/R204 samples (Fig 5C and D, S3 Fig).

RT-PCR assays have been developed to directly assess sub-genomic mRNA (sgRNA) as a measure of replicative intermediates of SARS-CoV-2 representing putative replication in cells rather than RNA packaged in virions or residual viral RNA (21, 22). A decline in sgRNA in sputum typically occurs from day 10 to 11 after onset of symptoms (22). Our finding that a variant can emerge that is associated with a novel sub-genomic RNA or may differentially impact the level of different sgRNAs suggest that the viral sequences should be analyzed to ensure the primers or probes used are appropriate and analysis of short read deep sequences with the periscope tool considered to help interpret results obtained from different variants.

### Potential impact of introduced TRS sequences on RNA structure

Modeling of the region around the mRNA encoding position 203 and 204 of the nucleocapsid using RNAfold (23) predicts the presence of a three-way junction in the RNA (S4 Fig), which was also predicted using Junction-Explorer (24). Three-way junction motifs are common throughout biology and are found both in pure RNAs, such as riboswitches or ribozymes, and in RNA-protein complexes, including the ribosome (25). RNA three-way junctions are often stabilized via terminal loop interactions with distant tertiary contacts while the junctions act like flexible hinges. These attributes allow these structures to sample unusual conformational spaces and they often form platforms for interactions with other molecules such as proteins, RNAs or small molecule ligands (25), and these folds often have an essential role in either the function or assembly of the molecules in which they are contained.

RNAfold predicts the mutation from A<u>GGG</u>GA to A<u>AAC</u>GA strongly disrupts this structure as the lengths of the predicted helices and each of the junctions are altered and the stability of Helix 2 is undermined (S4 Fig). A comparison of the two-modeled sequences using CHSalign (26) also indicates that none of the junctions are maintained. Given these widespread alterations, this modeling predicts that the A<u>GGG</u>GA to A<u>AAC</u>GA mutation would have a strong impact on the local RNA structure of this region, and likely impacts the normal function of this three-way junction motif. Interestingly, the RNA modeling shown in S4 Fig also suggests that pairing of specific nucleotides to maintain these RNA structures may require the preferential codon usage by RG (AGGGGA) and KR (AAACGA) and be a contributory factor to preferential codon usage in RNA viruses more generally even in protein coding regions.

While it is not possible to determine the impact of this proposed structural alteration on SARS-CoV-2 without a defined function for this structure, there are precedents where minor changes in a three-way junction have large functional consequences for their host viruses. For example, Flaviviruses such as Dengue and West Nile virus utilize the host cell machinery to degrade viral genomes until they encounter structures near the 3’ end that are resistant to XRN1 5’-3’ exonuclease (27). The resulting small flaviviral RNAs (sfRNAs) are non-coding RNAs that induce cytopathicity and pathogenicity. The resistance of sfRNA to XRN1 is dependent on the structure of a three-way junction and a single nucleotide change at the junction alters the fold sufficiently to prevent the accumulation of disease-related sfRNAs. Thus, small changes at the nucleotide level can have profound functional consequences for viral RNA three-way junctions.

### Lack of evidence that the RG to KR change at positions 203 and 204 of nucleocapsid was driven by HLA-restricted immune selective pressure

Selection of viral adaptations to polymorphic host responses mediated by T cells, NK-cells and antibodies are well described for other RNA viruses such as HIV and HCV (15, 28). HIV-1 adaptations to human leucocyte antigen (HLA)-restricted T-cell responses have also been shown to be transmitted and accumulate over time (29, 30). As previously shown for SARS-CoV, T-cell responses against SARS-CoV-2 are likely to target the nucleocapsid (31). Notably, SARS-CoV-2 R203K/G204R polymorphisms modify the predicted binding of putative HLA-restricted T-cell epitopes containing these residues (S2 Table). One of the predicted T-cell epitopes is restricted by the HLA-C*07 allele; and we therefore considered whether escape from HLA-C-restricted T-cell responses may conceivably confer a fitness advantage for SARS-CoV-2, particularly in European populations where HLA-C*07 is prevalent and carried by >40% of the population (www.allelefrequencies.net). However, using HLA-C*07:01 purified from the Steinlin cell line (IHWG ID: 9087; A*01:01, B*08:01 and C*07:01) and the anti-HLA Class I B123.2 mAb in inhibition assays we were not able to detect binding of either of the SARS-CoV-2 peptides S<u>RG</u>TSPARM or S<u>KR</u>TSPARM (S3 Table). We therefore have, as yet no evidence of any impact or selective advantage to the virus at the protein level of a change at position 203/204 from the RG to KR residues.

### SARS-CoV-2 and Host Adaptation: Implications for global viral dynamics, pathogenesis and immunogenicity

Currently the possible functional effect(s) of the introduction of the AAACGA motif from the leader TRS into the RNA encoding position 203 and 204 of the nucleocapsid at the RNA and protein level are not known. TRS sites are usually intergenic and it has been assumed that recombination events at such sites are more likely to be viable. It has also been shown recently that recombination breakpoint hotspots in coronaviruses are more frequently co-located with TRS-B sites than expected (32). Our findings suggest that a novel TRS-B site can be introduced in a recombination breakpoint from the leader TRS, and that this can occur within an ORF and remain viable. The exact mechanism by which the AAA CGA codons could have been incorporated from the TRS-L into the nucleocapsid is not known but may have first required the AAACGA to be captured from the TRS-L and then for replication to be reinitiated at the nucleocapsid to generate a full-length genomic RNA.

The nucleocapsid protein is a key structural protein critical to viral transcription and assembly (33), suggesting that changes in this protein could either increase or decrease replicative fitness. The K203/R204 polymorphism is located between the RNA binding/serine-rich domains and the dimerization structural domain (S5 Fig) in a part of the protein that has not been characterized in terms of 3-dimensional structure. The sequence of this region is not similar enough to solved structures to allow prediction of the influence of the K203/R204 polymorphisms on the structure or function of the protein. However, it is known that SARS-CoV-2 is exquisitely sensitive to interferons and that it depends on the nucleocapsid and M proteins to maintain interferon antagonism (34, 35). Specifically the C terminus (aa 362 to 422) of the nucleocapsid, which is predicted to be expressed at higher levels in those with the KR variant and novel sgRNA, has been shown to interact with the SPRY domain of TRIM25 disturbing its interaction with CARDs of RIG-I inhibiting RIG-I ubiquitination and Type 1 interferon signaling (36). Importantly the cells expressing the C-terminal nucleocapsid protein in that study produced lower viral titer, suggesting the incorporation of this protein into the nucleocapsid may reduce the formation of functional virus. This raises the possibility that any enhancement of inhibition of interferon signaling associated with the novel K203/R204 sgRNA may be offset by less efficient replication, potentially accounting for the lack of association with higher viral load in the upper respiratory tract and absence of epidemiologic evidence of increased transmission. It is also possible that the increase in sgRNA directly inhibits RIG-I signaling and downstream Type I interferon responses as has been described for Dengue serotype 2 (37). Finally, the central region of coronavirus nucleocapsid (aa 117 to 268) has been shown to have RNA chaperone activity that enhances template switching and efficient transcription possibly accounting for the increase in sgRNA for the E and M proteins and ORF6 in KR-sequences compared to RG-sequences (38). Note we cannot exclude that the novel sgRNA may also use the downstream ATG in the ORF9c reading frame.

The adaptive potential of differential expression of sgRNAs is supported by a recent study by Thorne and colleagues that demonstrates that the B.1.1.7 (‘Alpha’ or UK variant) isolate containing the R203K/G204R substitutions is associated with enhanced antagonism of the innate immune response (39). Specifically, this study showed that in-vitro infection of human lung epithelial (Calu-3) cells by B.1.1.7 isolates showed diminished RNA and protein expression of IFNβ and reduced induction of interferon sensitive genes relative to other isolates without these defining mutations in the nucleocapsid (normalized for intracellular viral RNA). This effect was independent of the reduced sensitivity to type I and III IFNs described for isolates carrying the D614G spike mutation (40). Further evaluation of this system showed that infection with the B.1.1.7 isolate resulted in significant changes in protein expression of known innate immune regulators such as ORF9b (41), ORF6 (42) and nucleocapsid (36, 43), as well as increased levels of the N* sgRNA described in this study and was again confirmed to be unique to those isolates with the R203K/G204R mutations. These increased levels of sgRNAs and protein support the findings in this study showing increased sgRNA levels for N, ORF6 and N* in clinical samples from B.1.1.7-infected subjects relative to subjects infected with other SARS-CoV-2 isolates. Interestingly, the increased levels of ORF9b may be due to the D3L mutation in the nucleocapsid that we have proposed to have arisen similarly to the R203G/G204R mutations and is associated with increased levels of B.1.1.7 sgRNA encoding ORF9b in clinical samples (44).

The B.1.617.2 (‘Delta’) variant appears to be more transmissible even in the context of previous vaccination and is now replacing other variants. This variant has acquired an R203M substitution as a result of a single nucleotide change while retaining an arginine (G) at position 204. This raises the possibility that the residue 203 is critical to the interaction of nucleocapsid with TRIM25 decreasing the Type 1 interferon response or increases transmissibility in some other way (36).

Other contemporary concerns include the fall in antibody levels following infection or vaccination, the potential limited durability of protection afforded by currently available vaccines and the risk of reinfection by variants after vaccination (45, 46). At low levels of antibodies, the lungs appear to remain relatively protected against severe disease presumably by some combination of antibodies and amnestic responses restimulated by the time the lung is involved. In contrast, the early establishment of infection in the upper respiratory tract appears possible if antibody levels are low (47). We therefore postulate that variants that are more effective in interfering with Type 1 interferon responses would be more transmissible, but not necessarily cause severe disease in the context of waning immunity at an individual or population level.

## Conclusion

Marked viral diversity and adaptation of other RNA viruses such as HIV, HCV and influenza to host selective pressures have been a barrier to successful treatment and vaccination to date. Although SARS-CoV-2 is less diverse and adaptable, the D614G variant and the K203/R204 and Delta variants have emerged by either nucleotide mutation or homologous recombination during its rapid, widespread global spread and do appear to have functional impact. It will therefore be critical to continue molecular surveillance of the virus and elucidate the functional consequences of any newly emerging viral genetic changes to guide development of diagnostics, antivirals and universal vaccines and to target conserved and potentially less mutable SARS-CoV-2 elements. The ability of SARS-CoV-2 to introduce new TRS motifs throughout its genome with the potential to introduce both novel sub-genomic RNA transcripts and coding changes in its proteins may add to these challenges.

## Acknowledgments

We thank colleagues at the Institute for Immunology and Infectious Diseases, Murdoch University, Australia and the Department of Medicine, Division of Infectious Diseases, Vanderbilt University Medical Center, USA. We would like to acknowledge additional members of the Sheffield COVID-19 Genomic Group who contributed to the generation of the sequence data: Adrienne Angyal, Rebecca L. Brown, Laura Carrilero, Cariad M Evans, Luke R. Green, Danielle C. Groves, Katie J Johnson, Paul J Parsons, David Partridge, Mohammad Raza, Rachel M. Tucker, Dennis Wang, Matthew D. Wyles. Ethics approval and consent to participate (COG-UK CONSORTIUM; R&D NR0195).

We thank the reviewers and editor for their helpful comments.

