## Supplementary Material for "Generation of a novel SARS-CoV-2 sub-genomic RNA due to the R203K/G204R variant in nucleocapsid: homologous recombination has potential to change SARS-CoV-2 at both protein and RNA level"

**This file includes:**

Figures S1 to S5

Tables S1 to S9

**Accession numbers:**

Metatranscriptome data from coronaviruses in acute respiratory infections and asymptomatic subjects:

| | | | |
|---|---|---|---|
| Coronavirus_NL63_S168.sqn | PRJNA671738 | SAMN16547776 | SRR12893437 |
| Coronavirus_NL63_S170.sqn | PRJNA671738 | SAMN16547777 | SRR12893436 |
| Coronavirus_OC43_S219.sqn | PRJNA671738 | SAMN16547778 | SRR12893435 |
| Coronavirus_229E_S220.sqn | PRJNA671738 | SAMN16547779 | SRR12893434 |

Data for clinical cohort at https://www.cogconsortium.uk/data/.

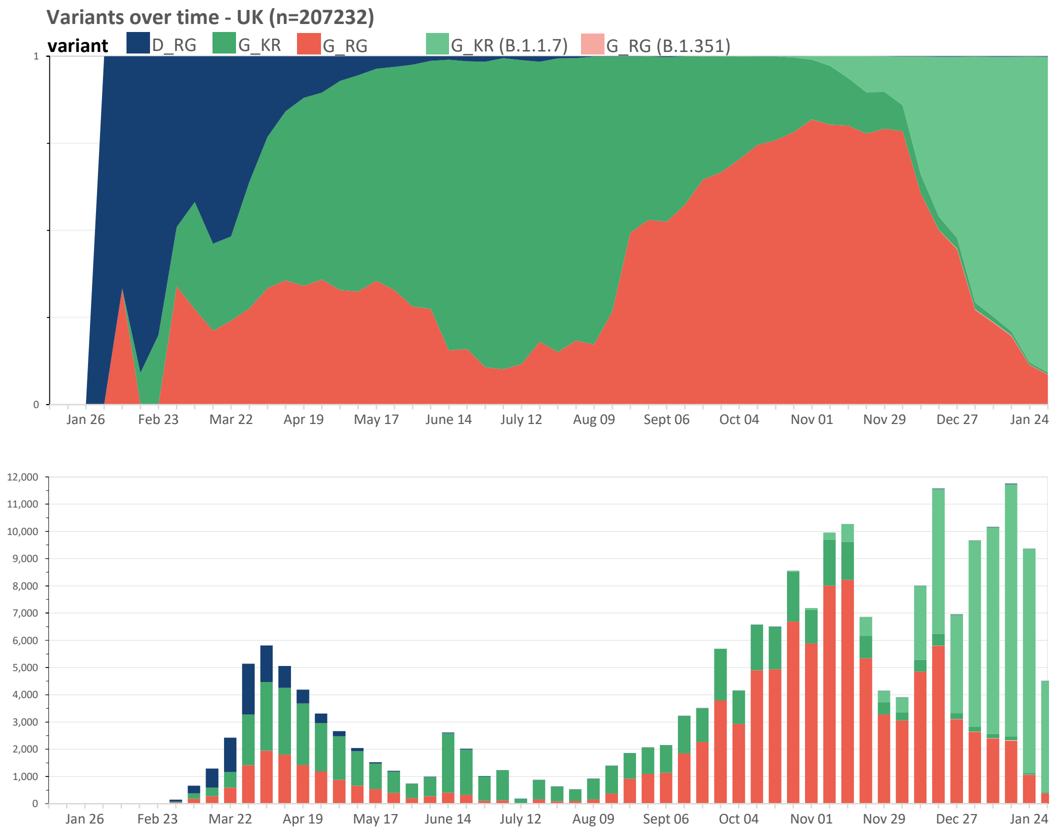

**S1 Fig. Proportion of weekly deposited SARS-CoV-2 sequences in the UK.** Top panel is the proportion of different strains. Bottom panel is the number of deposited sequences in the GISAID database broken down into specific variants. The B.1.1.7 'UK variant' is the main deposited strain in recent months. D_RG = D614/R203/G204; G_RG = G614/R203/G204; G_KR = G614/K203/R204; G_KR (B.1.1.7) = 'UK variant'; and G_RG (B.1.351) = 'South African variant'.

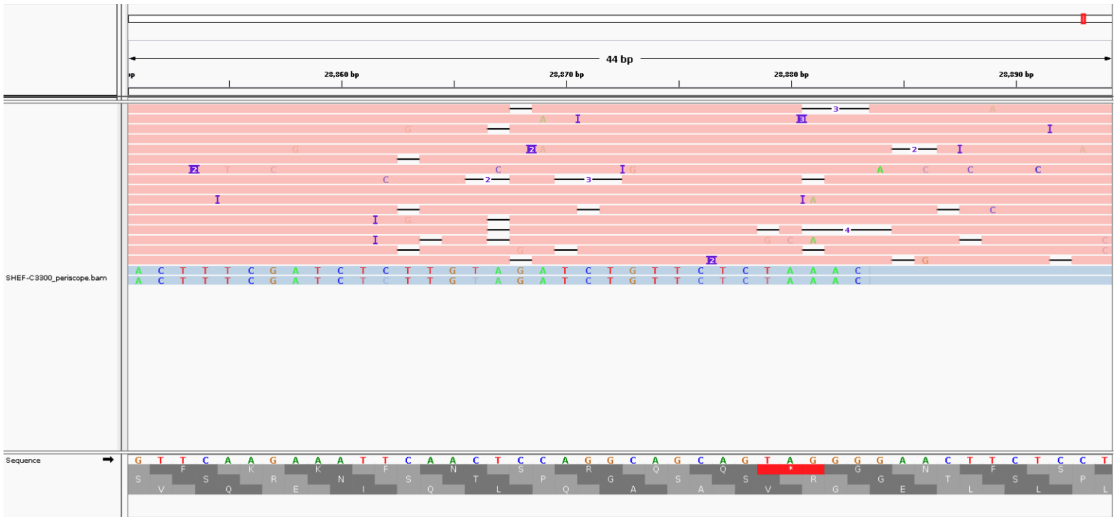

**S2 Fig. Possible erroneous demultiplexing is responsible for novel sgRNA detected in a single sequence with R203/G204.** This sample does not have the R203K/G204R mutation but two reads were classified as novel sgRNA by the periscope tool (blue). Further investigation shows that this sample, indeed, does not contain any evidence of the new TRS (red reads). It is possible that these reads are due to a barcoding issue or a very low level of sample contamination.

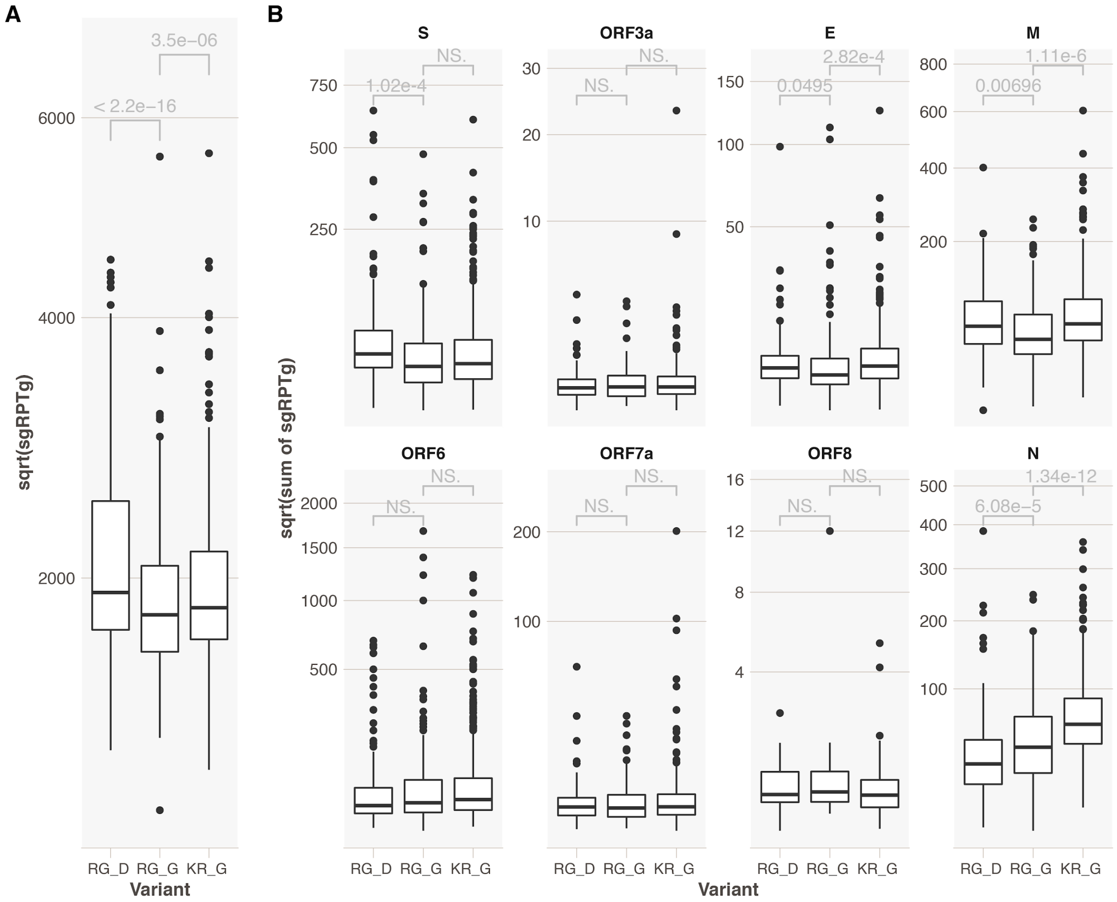

**S3 Fig. Sub-genomic RNA levels considering the D614 site in spike protein.** Samples with a D at residue 614 in the spike protein appear to have increased expression of sgRNA compared to those with a G at this residue. (A). Across all sgRNAs and (B) in selected individual sgRNAs (S = spike, M = membrane and E = envelope). Y-axis is square root transformed sgRNA reads per 1000 genomic RNA reads from corresponding amplicons. p values from Mann-Whitney U, adjusted for multiple testing with the Holm method. RG = R203/G204 containing variant; KR = K203/R204 containing variants; D = spike D614 containing variants, G = spike G614 containing variants.

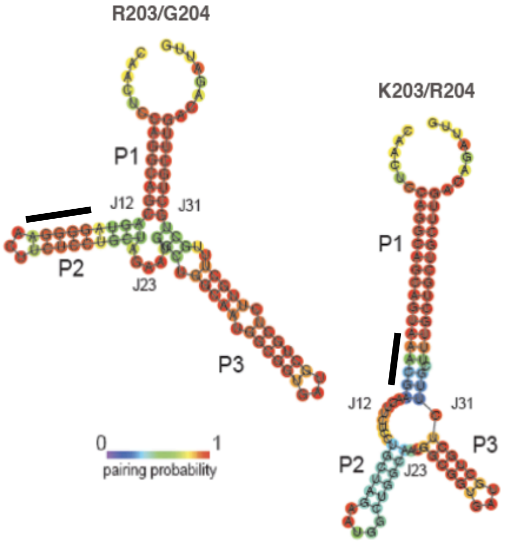

**S4 Fig. Predicted RNA structures corresponding to the R203/G204 and K203/R204 forms of the nucleocapsid suggest alterations to the three-way junctions.** Predictions performed using the RNAfold program with pairing probability shown. Black bars highlight the 203/204 codons.

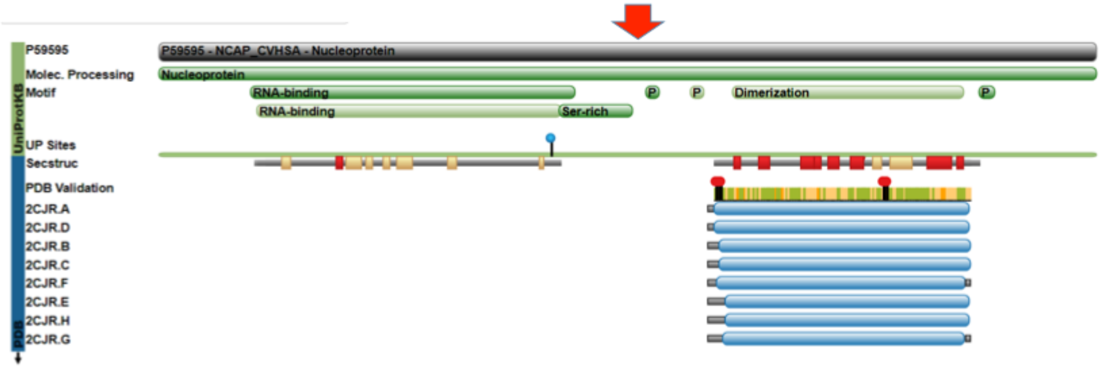

**S5 Fig. Location of R203K/G204R polymorphisms between the RNA-binding and dimerization domains.** The grey horizontal bar indicates the full length nucleocapsid protein with the individual domains and specific regions indicated by green horizontal bars. The left side panel indicates the presence of structures in the public databases. There are no structures for the region adjacent to the serine-rich stretch and containing the R203K/G204R sites (indicated by a red arrow). Structures are available for the RNA-binding domain (including for SARS-CoV-2) and dimerization domain (SARS-CoV only).

**S1 Table. Amino acid variations and alternative codon usage of SARS-CoV-2 (>5% frequency of deposited sequences; 24th January 2021).**

| Gene / Protein [Length] Amino Acid Position | Codon Usage | | |
|---|---|---|---|
| ORF1ab / ORF1ab protein [7097] | Amino Acid [Codon Count] (5% cutoff) | | |
| 60 | V [GTT 78] | [GTC 21] | |
| 216 | S [TCC 88] | [TCT 11] | |
| 265 | T [ACC 85] | | I [ATC 14] |
| 924 | F [TTT 94] | [TTC 5] | |
| 1001 | T [ACT 88] | | I [ATT 11] |
| 1708 | A [GCT 88] | | D [GAT 11] |
| 1907 | F [TTC 87] | [TTT 12] | |
| 2007 | T [ACC 78] | [ACT 21] | |
| 2230 | I [ATA 88] | | T [ACA 11] |
| 3606 | L [TTG 94] | | F [TTT 5] |
| 4715 | L [CTT 94] | | P [CCT 5] |
| 4804 | P [CCC 88] | [CCT 11] | |
| 5005 | H [CAC 88] | [CAT 11] | |
| 5304 | T [ACT 88] | [ACC 11] | |
| 6205 | L [CTA 94] | [TTA 5] | |
| 6668 | L [TTA 93] | [TTG 6] | |
| 6997 | A [GCG 77] | [GCC 22] | |

| S / surface protein [1274] | Amino Acid [Codon Count] (5% cutoff) | |
|---|---|---|
| 18 | L [CTT 90] | F [TTT 9] |
| 222 | A [GCT 77] | V [GTT 22] |
| 477 | S [AGC 94] | N [AAC 5] |
| 501 | N [AAT 87] | Y [TAT 12] |
| 570 | A [GCT 88] | D [GAT 11] |
| 614 | G [GGT 94] | D [GAT 5] |
| 681 | P [CCT 87] | H [CAT 12] |
| 716 | T [ACA 88] | I [ATA 11] |
| 982 | S [TCA 88] | A [GCA 11] |
| 1118 | D [GAC 88] | H [CAC 11] |

| ORF3a [276] | Amino Acid [Codon Count] (5% cutoff) | |
|---|---|---|
| 57 | Q [CAG 78] | H [CAT 21] |

| M / membrane glycoprotein [223] | Amino Acid [Codon Count] (5% cutoff) | |
|---|---|---|
| 71 | Y [TAC 94] | [TAT 5] |
| 93 | L [CTC 77] | [CTG 21] |

| ORF8 / OFR8 protein [122] | Amino Acid [Codon Count] (5% cutoff) | | |
|---|---|---|---|
| 17 | H [CAC 85] | [CAT 15] | |
| 24 | S [TCA 94] | | L [TTA 5] |
| 27 | Q [CAA 88] | | * [TAA 11] |
| 52 | R [AGA 88] | | I [ATA 11] |

| | | |
|---|---|---|
| **73** | Y [TAC 88] | C [TGC 11] |

| **N / nucleocapsid [421]** | **Amino Acid [Codon Count] (5% cutoff)** | |
|---|---|---|
| **3** | D [GAT 88] | L [CTA 11] |
| **194** | S [TCA 94] | L [TTA 5] |
| **199** | P [CCA 94] | L [CTA 5] |
| **203** | R [AGG 62] | K [AAA 37] |
| **204** | G [GGA 62] | R [CGA 37] |
| **220** | A [GCT 78] | V [GTT 21] |
| **235** | S [TCT 88] | F [TTT 11] |

| **ORF10 [276]** | **Amino Acid [Codon Count] (5% cutoff)** | |
|---|---|---|
| **30** | V [GTA 78] | L [TTA 21] |

**S2 Table. Peptide prediction for regions in SARS-CoV-2 containing the R203K/G204R amino acid combinations.***

| NetMHC | HLA | Peptide | 1-log50k(aff) | Affinity(nM) | %Rank | Bind Level |
|---|---|---|---|---|---|---|
| KR | HLA-A 30:01 | **R**TSPARMAG | 0.603 | 73.42 | 0.5 | SB |
| RG | HLA-A 30:01 | **G**TSPARMAG | 0.342 | 1235.69 | 2.5 | |
| KR | HLA-A 68:01 | NSTPGSS**KR** | 0.666 | 37.12 | 0.5 | SB |
| RG | HLA-A 68:01 | NSTPGSS**RG** | 0.063 | 25298.77 | 22 | |
| KR | HLA-B 15:03 | S**KR**TSPARM | 0.725 | 19.68 | 0.3 | SB |
| RG | HLA-B 15:03 | S**RG**TSPARM | 0.273 | 2606.69 | 6 | |
| KR | HLA-B 73:01 | **KR**TSPARMA | 0.166 | 8306.36 | 0.5 | SB |
| RG | HLA-B 73:01 | **RG**TSPARMA | 0.035 | 34081.84 | 11 | |
| KR | HLA-C 07:01 | S**KR**TSPARM | 0.104 | 16237.59 | 6.5 | |
| RG | HLA-C 07:01 | S**RG**TSPARM | 0.248 | 3406.51 | 1.2 | WB |
| KR | HLA- C07:02 | S**KR**TSPARM | 0.216 | 4831.5 | 1.5 | WB |
| RG | HLA- C07:02 | S**RG**TSPARM | 0.296 | 2035.83 | 0.7 | WB |

| NetMHCpan | HLA | Peptide | Score | %Rank | Bind Level |
|---|---|---|---|---|---|
| KR | HLA-A 30:01 | RNSTPGSS**K** | 0.281867 | 0.4783 | SB |
| RG | HLA-A 30:01 | RNSTPGSS**R** | 0.092325 | 2.6775 | |
| KR | HLA-A 30:01 | **R**TSPARMAG | 0.438433 | 0.1607 | SB |
| RG | HLA-A 30:01 | **G**TSPARMAG | 0.162082 | 1.3184 | WB |
| KR | HLA-A 68:01 | NSTPGSS**KR** | 0.705615 | 0.1942 | SB |
| RG | HLA-A 68:01 | NSTPGSS**RG** | 0.00035 | 35.9355 | |
| KR | HLA-A 33:03 | NSTPGSS**KR** | 0.378816 | 0.3332 | SB |
| RG | HLA-A 33:03 | NSTPGSS**RG** | 0.000122 | 54.0541 | |
| KR | HLA-C 07:01 | S**KR**TSPARM | 0.041824 | 3.6073 | |
| RG | HLA-C 07:01 | S**RG**TSPARM | 0.318753 | 0.3051 | SB |
| KR | HLA-C 07:02 | S**KR**TSPARM | 0.059982 | 4.5638 | |
| RG | HLA-C 07:02 | S**RG**TSPARM | 0.355166 | 0.5579 | WB |
| KR | HLA-B 38:01 | S**KR**TSPARM | 0.003955 | 13.1727 | |
| RG | HLA-B 38:01 | S**RG**TSPARM | 0.080137 | 1.8794 | WB |
| KR | HLA-B 14:02 | S**KR**TSPARM | 0.043487 | 3.2691 | |
| RG | HLA-B 14:02 | S**RG**TSPARM | 0.080815 | 1.7515 | WB |

**SB = strong binder and WB = weak binder. Sites 203 and 204 indicated in red.*

**S3 Table. Binding affinity of peptides to specific HLA alleles including peptides containing the R203K/G204R variants.**

| | | | | | Binding affinity (IC50 nM) | |
|---|---|---|---|---|---|---|
| Peptide ID | Sequence | Len | Source | Notes | B*08:01 | C*07:01 |
| 1054.0002 | FLRGRAYGI | 9 | HSV nuc 11 | HLA B8 T cell epitope | **0.85** | - |
| 4199.0002 | QAKWRLQTL | 9 | HSV nuc 26 | B08 tetramer; PMID:11927633 | **2.4** | - |
| 4199.0001 | EIYKRWII | 8 | HIV gag 260 | B08 tetramer; PMID:27760342 | **2.6** | - |
| 960.0002 | FLKDYQLL | 8 | HIV gp 586 | Analog of B8 epitope | **6.9** | - |
| 3484.0028 | IRSSYIRVL | 9 | Mamu DNA rep factor 289 | Mamu B*1001/HLA C*07:01 ligand | **301** | **0.21** |
| 4196.0001 | YQSGLSIVM | 9 | MTB hyp protein 48 | C*0701 binder; PMID:23555576 | 1107 | **67** |
| 4196.0002 | ANNTRLWVY | 9 | MTB ag 85B | C*0701 tetramer; PMID:25809751 | 42748 | 1600 |
| 1074.0001 | YTAVVPLVY | 9 | Hu J chain 102 | HLA A1 eluted ligand | 29285 | 1279 |
| 4197.0001 | SRGTSPARM | 9 | SARS-CoV-2 nuc 202 | | - | - |
| 4197.0002 | SKRTSPARM | 9 | SARS-CoV-2 nuc 202 | | 10620 | - |

A dash indicates IC50 >50000 nM.

**S4 Table. Frequency of linked amino acid variations across the SARS-CoV-2 genome (>0.01% frequency of deposited sequences; 24th January 2021).***

| N | | ORF1ab | | | | | | | | | S | | | | | | | | | | | | | ORF3 | ORF8 | | | | N | | | | | ORF10 | |
|---|---|---|---|---|---|---|---|---|---|---|---|---|---|---|---|---|---|---|---|---|---|---|---|---|---|---|---|---|---|---|---|---|---|---|---|
| KR# | RG# | T 265 I | T 1001 I | A 1708 D | I 2230 T | L 3606 F | SGF 3675-3577 deletion | | | L 4715 P | L 18 F | HV 69-70 deletion | | Y 144 deletion | A 222 V | S 477 N | N 501 Y | A 570 D | G 614 D | P 681 H | T 716 I | S 982 A | D 1118 H | Q 57 H | S 24 L | Q 27 * | R 52 I | Y 73 C | D 3 L | S 194 L | P 199 L | A 220 V | S 235 F | V 30 L | |
| | | 1058 | 3266 | 5387 | 6953 | 11081 | 11288 | 11291 | 11294 | 14407 | 21614 | 21767 | 21770 | 21992 | 22226 | 22991 | 23063 | 23270 | 23402 | 23603 | 23708 | 24506 | 24914 | 25561 | 27963 | 27972 | 28047 | 28110 | 28280 | 28853 | 28868 | 28931 | 28976 | 29645 | 28880 - 28885 |
| 0 | 11834 | T | T | A | I | L | S | G | F | P | L | H | V | Y | A | S | N | A | D | P | T | S | D | Q | S | Q | R | Y | D | S | P | A | S | V | WILDTYPE |
| 88812 | 27318 | T | T | A | I | L | S | G | F | L | L | H | V | Y | A | S | N | A | G | P | T | S | D | Q | S | Q | R | Y | D | S | P | A | S | V | |
| 50179 | 0 | T | I | D | T | L | - | - | - | L | L | - | - | - | A | S | Y | D | G | H | I | A | H | Q | S | * | I | C | L | S | P | A | F | V | |
| 9670 | 620 | T | T | A | I | L | S | G | F | L | L | H | V | Y | A | N | N | A | G | P | T | S | D | Q | S | Q | R | Y | D | S | P | A | S | V | |
| 5370 | 848 | T | T | A | I | F | S | G | F | L | L | H | V | Y | A | S | N | A | G | P | T | S | D | Q | S | Q | R | Y | D | S | P | A | S | V | |
| 1174 | 0 | T | I | D | T | L | - | - | - | L | F | - | - | - | A | S | Y | D | G | H | I | A | H | Q | S | * | I | C | L | S | P | A | F | V | |
| 783 | 87 | T | T | A | I | L | S | G | F | L | L | H | V | Y | A | S | N | A | G | H | T | S | D | Q | S | Q | R | Y | D | S | P | A | S | V | |
| 720 | 4 | T | T | A | I | L | S | G | F | L | L | - | - | Y | A | S | N | A | G | P | T | S | D | Q | S | Q | R | Y | D | L | P | A | S | V | |
| 690 | 13 | T | T | A | I | L | S | G | F | L | L | H | V | Y | A | S | N | A | G | P | T | S | D | Q | S | Q | R | Y | D | S | S | A | S | V | |
| 636 | 1 | T | T | A | I | L | S | G | F | L | L | H | V | Y | A | S | Y | A | G | P | T | S | D | Q | S | Q | R | Y | D | S | P | A | S | V | |
| 401 | 12 | T | T | A | I | L | S | G | F | L | L | H | V | Y | V | S | N | A | G | P | T | S | D | Q | S | Q | R | Y | D | S | P | A | S | V | |
| 362 | 0 | T | T | A | I | L | S | G | F | L | L | H | V | Y | A | R | N | A | G | P | T | S | D | Q | S | Q | R | Y | D | S | P | A | S | V | |
| 342 | 0 | T | I | D | T | F | - | - | - | L | L | - | - | - | A | S | Y | D | G | H | I | A | H | Q | S | * | I | C | L | S | P | A | F | V | |
| 232 | 17 | T | T | A | I | L | S | G | F | L | L | H | V | Y | A | S | N | A | G | P | T | S | D | Q | S | Q | R | Y | D | S | P | A | S | L | |
| 223 | 99 | T | T | A | I | L | S | G | F | L | F | H | V | Y | A | S | N | A | G | P | T | S | D | Q | S | Q | R | Y | D | S | P | A | S | V | |
| 201 | 9723 | T | T | A | I | L | S | G | F | L | L | H | V | Y | A | S | N | A | G | P | T | S | D | H | S | Q | R | Y | D | S | P | A | S | V | |
| 165 | 11438 | T | T | A | I | L | S | G | F | L | L | H | V | Y | A | S | N | A | G | P | T | S | D | Q | S | Q | R | Y | D | L | P | A | S | V | |
| 156 | 82 | T | T | A | I | L | S | G | F | L | L | H | V | - | A | S | N | A | G | P | T | S | D | Q | S | Q | R | Y | D | S | P | A | S | V | |
| 148 | 5 | T | I | A | I | L | S | G | F | L | L | H | V | Y | A | S | N | A | G | P | T | S | D | Q | S | Q | R | Y | D | S | P | A | S | V | |
| 129 | 33 | T | T | A | I | L | S | G | F | L | L | H | V | Y | A | S | N | A | G | P | T | S | D | Q | S | * | R | Y | D | S | P | A | S | V | |
| 121 | 0 | T | T | A | I | L | S | G | F | L | L | H | V | Y | A | I | N | A | G | P | T | S | D | Q | S | Q | R | Y | D | S | P | A | S | V | |
| 114 | 5 | T | T | A | I | L | S | G | F | L | L | H | V | Y | A | S | N | A | G | R | T | S | D | Q | S | Q | R | Y | D | S | P | A | S | V | |
| 113 | 0 | T | T | A | I | F | S | G | F | L | L | - | - | Y | A | S | N | A | G | P | T | S | D | Q | S | Q | R | Y | D | L | P | A | S | V | |
| 113 | 1 | T | T | A | I | L | F | G | F | L | L | H | V | Y | A | S | N | A | G | P | T | S | D | Q | S | Q | R | Y | D | S | P | A | S | V | |
| 112 | 0 | T | I | D | T | L | S | G | L | L | L | - | - | - | A | S | Y | D | G | H | I | A | H | Q | S | * | I | C | L | S | P | A | F | V | |
| 103 | 237 | T | T | A | I | L | S | G | F | P | L | H | V | Y | A | S | N | A | G | P | T | S | D | Q | S | Q | R | Y | D | S | P | A | S | V | |
| 92 | 1 | T | T | A | I | F | S | G | F | L | L | H | V | Y | A | N | N | A | G | P | T | S | D | Q | S | Q | R | Y | D | S | P | A | S | V | |
| 88 | 12 | T | T | A | I | L | S | G | F | L | L | H | V | Y | A | S | N | A | G | P | T | S | D | Q | S | Q | R | Y | Y | S | P | A | S | V | |
| 87 | 106 | I | T | A | I | L | S | G | F | L | L | H | V | Y | A | S | N | A | G | P | T | S | D | Q | S | Q | R | Y | D | S | P | A | S | V | |
| 85 | 0 | T | T | A | I | L | - | - | - | L | F | H | V | Y | A | S | Y | A | G | P | T | S | D | Q | S | Q | R | Y | D | S | P | A | S | V | |
| 85 | 65 | T | T | A | I | L | S | G | F | L | L | H | V | Y | A | S | N | A | D | P | T | S | D | Q | S | Q | R | Y | D | S | P | A | S | V | |
| 81 | 17 | T | T | A | I | L | S | G | F | L | L | H | V | Y | A | S | N | A | G | P | I | S | D | Q | S | Q | R | Y | D | S | P | A | S | V | |
| 78 | 6 | T | T | A | I | L | S | G | F | L | L | H | V | Y | A | S | N | A | G | L | T | S | D | Q | S | Q | R | Y | D | S | P | A | S | V | |
| 58 | 6750 | T | T | A | I | L | S | G | F | L | L | - | - | Y | A | S | N | A | G | P | T | S | D | Q | S | Q | R | Y | D | S | P | A | S | V | |
| 57 | 1 | T | T | A | I | L | S | G | F | L | L | H | F | Y | A | S | N | A | G | P | T | S | D | Q | S | Q | R | Y | D | S | P | A | S | V | |
| 54 | 0 | T | T | A | I | L | S | G | F | L | L | H | V | Y | A | S | N | A | G | P | T | S | Y | Q | S | Q | R | Y | D | S | P | A | S | V | |
| 34 | 32622 | I | T | A | I | L | S | G | F | L | L | H | V | Y | A | S | N | A | G | P | T | S | D | H | S | Q | R | Y | D | S | P | A | S | V | |
| 17 | 11834 | T | T | A | I | L | S | G | F | P | L | H | V | Y | A | S | N | A | D | P | T | S | D | Q | S | Q | R | Y | D | S | P | A | S | V | |
| 15 | 5434 | T | T | A | I | L | S | G | F | L | L | H | V | Y | A | S | N | A | G | P | T | S | D | Q | S | Q | R | Y | D | S | L | A | S | V | |
| KP 208 | | T | I | D | T | L | - | - | - | L | L | - | - | - | A | S | Y | D | G | H | I | A | H | Q | S | * | I | C | L | S | P | A | F | V | |

*Highlighted are the defining variations of the B.1.1.7 variant. Note the K203/P204 variant shown at the bottom is likely to have arisen from the B.1.1.7 UK variant.*

**S5 Table. Variants R203/G204 and K203/R204 are the main amino acid combinations at positions 203 and 204 in nucleocapsid*.**

| 203/204 amino acid | Count [codon combination] | | % deposited sequences |
|---|---|---|---|
| RG | 302305 [AGG] [GGA] | 31 [AGA] [GGA] | 62.2 |
| KR | 181752 [AAA] [CGA] | | 37.4 |
| KL | 915 [AAA] [CTA] | | 0.19 |
| KG | 414 [AAG] [GGA] | | 0.1 |
| KP | 240 [AAA] [CCA] | | <0.1 |
| MG | 207 [ATG] [GGA] | | <0.1 |
| RR | 98 [AGG] [AGA] | | <0.1 |
| SG | 93 [AGT] [GGA] | | <0.1 |

**Global SARS-CoV-2 sequences with sequence coverage of nucleocapsid amino acid positions 203 and 204 downloaded from www.gisaid.org on 24th of January 2021.*

**S6 Table. Frequency of sgRNA transcripts in 90 individuals that carry either the K203/R204 or R203/G204 variant from the SRA database (www.ncbi.nlm.nih/sra).***

| 45 KR Samples (leader sequence counts) | | | Sample Count 2 MMs | |
|---|---|---|---|---|
| Leader Start Base Position | TRS Start Base Position | Gene / Protein | Sample Count | Read Count |
| 39 | 66 | ORF1a | 45 | 10087 |
| 3784 | 3811 | | 1 | 1 |
| 5675 | 5702 | | 1 | 2 |
| 10609 | 10636 | | 1 | 1 |
| 14340 | 14367 | | 1 | 1 |
| 18910 | 18937 | | 1 | 1 |
| 21525 | 21552 | S | 17 | 125 |
| 22116 | 22143 | | 1 | 1 |
| 23822 | 23849 | | 2 | 2 |
| 25354 | 25381 | ORF3a | 9 | 13 |
| 26202-26208 | 26229-26235 | E | 19 | 93 |
| 26442 | 26469 | M | 13 | 44 |
| 27010 | 27037 | ORF6 | 21 | 37 |
| 27356-27359 | 27383-27386 | ORF7a | 14 | 22 |
| 27643 | 27670 | | 1 | 1 |
| 27857 | 27884 | ORF8 | 1 | 2 |
| 28134 | 28161 | | 1 | 1 |
| 28228-28233 | 28255-28260 | N | 40 | 704 |
| 28851 | 28878 | N KR | 5 | 6 |

| TRS sequence counts | | |
|---|---|---|
| ORF1a | 45 | 57989 |
| S | 45 | 49243 |
| ORF3a | 40 | 150 |
| E | 33 | 380 |
| M | 45 | 114549 |
| ORF6 | 31 | 187 |
| ORF7a | 39 | 306 |
| ORF8 | 45 | 221224 |
| N | 45 | 288450 |
| N KR | 27 | 61 |

| 45 RG Samples (leader sequence counts) | | | Sample Count 2 MMs | |
|---|---|---|---|---|
| Leader Start Base Position | TRS Start Base Position | Gene / Protein | Per Sample | Read Count |
| 39 | 66 | ORF1a | 45 | 14261 |
| 462 | 489 | | 1 | 1 |
| 3784 | 3811 | | 1 | 1 |
| 3886 | 3913 | | 1 | 1 |
| 5561 | 5588 | | 1 | 2 |
| 9562 | 9589 | | 1 | 1 |
| 12938 | 12965 | | 1 | 2 |
| 19081 | 19108 | | 1 | 1 |
| 21040 | 21067 | | 1 | 1 |
| 21525 | 21552 | S | 28 | 81 |
| 22470 | 22497 | | 2 | 2 |
| 22528 | 22555 | | 1 | 2 |
| 25364 | 25381 | ORF3a | 6 | 10 |
| 25391 | 25418 | | 1 | 1 |
| 25633 | 25660 | | 1 | 6 |
| 26206 | 26233 | E | 24 | 53 |
| 26260 | 26287 | | 1 | 1 |
| 26442 | 26469 | M | 22 | 64 |
| 27010 | 27037 | ORF6 | 22 | 44 |
| 27367 | 27384 | ORF7a | 6 | 10 |
| 27857 | 27884 | ORF8 | 2 | 3 |
| 28228-28233 | 28255-28260 | N | 41 | 342 |
| 29360 | 29377 | | 1 | 2 |

| TRS sequence counts | | |
|---|---|---|
| ORF1a | 45 | 85214 |
| S | 45 | 44171 |
| ORF3a | 30 | 101 |
| E | 37 | 311 |
| M | 45 | 130459 |
| ORF6 | 39 | 288 |
| ORF7a | 38 | 206 |
| ORF8 | 45 | 239659 |
| N | 45 | 326435 |
| N RG | 0 | 0 |

**The top two tables represent matches spanning position 6 to 27 of the leader sequence with up to two mismatches. The bottom two tables represent a relaxation of the criterion for partial leader sequence matches to allow for the known poor quality sequence at the 5' end of sequence reads. Highlighted in red is the novel non-canonical nucleocapsid sgRNA.*

**S7 Table. Risk of admission to critical care unit according to age, sex and R203/G204 vs K203/R204 status of infecting SARS-CoV-2 strain.***

| | **Odds Ratio** | **95% CI** | **P value** |
|---|---|---|---|
| Age in years | 1.00 | 0.99 - 1.02 | 0.898 |
| Sex (Male) | 4.16 | 2.09 - 8.88 | 9.43E-05 |
| K203/R204 | 1.20 | 0.63 - 2.34 | 0.588 |

**Multivariable logistic regression model using data from 981 individuals sampled in Sheffield, UK.*

**S8 Table. Impact of extraction method, day of illness at sampling and spike 614/nucleocapsid 203/204 variant on E gene cycle threshold (CT) value (A) G_RG and G_KR estimates using D_RG as reference and (B) D_RG and G_KR estimates using G_RG as reference.***

**A**

| | | Estimate | 95% CI | P value |
|---|---|---|---|---|
| Extraction_method (heat inactivation) | | 3.76 | 2.94 – 4.58 | <2.00E-16 |
| Days from symptom onset | | 0.42 | 0.23 – 0.61 | 2.05E-05 |
| Spike 614 & Nucleocapsid 203/204 status: **Reference D_RG (wild type)** | G_RG | -2.01 | -3.12 to -0.68 | 0.00011 |
| | G_KR | -1.90 | -3.01 to -1.00 | 0.0023 |

**B**

| | | Estimate | 95% CI | P value |
|---|---|---|---|---|
| Extraction_method (heat inactivation) | | 3.76 | 2.94 – 4.58 | <2.00E-16 |
| Days from symptom onset | | 0.42 | 0.23 – 0.61 | 2.05E-05 |
| Spike 614 & Nucleocapsid 203/204 status: **Reference G_RG** | D_RG | 1.90 | 0.68 – 3.12 | 0.0023 |
| | G_KR | -0.16 | -1.07 to 0.86 | 0.83 |

**Results from multivariable linear regression models. n=478 individuals sampled in Sheffield, UK). D_RG = D614/R203/G204; G_RG = G614/R203/G204; G_KR = G614/K203/R204. As due to reagent availability, method of extraction from clinical diagnostic samples changed during the study from the Magnapure96-based extraction to heat inactivation alone, this variable was included in the models. Heat inactivation (compared to Magnapure96 extraction) and later day from symptom onset were both associated with higher CT values (lower viral loads). K203/204 status is not associated with a change in CT value (A), whereas D614G status is associated with lower CT values/high viral loads (B). Of note K203/R204 samples form a 'subset' of D614G-containing variants.*

**S9 Table. Impact day of illness at sampling and spike 614/nucleocapsid 203/204 variant on total canonical sub-genomic RNA levels (A) G_RG and G_KR estimates using D_RG as reference and (B) D_RG and G_KR estimates using G_RG as reference.***

**A**

| **sgRNA expression** | | **Estimate** | **95% CI** | **P value** |
|---|---|---|---|---|
| Days from symptom onset | | 0.61 | 0.39 – 0.84 | 9.9E-08 |
| Spike 614 & Nucleocapsid 203/204 status: **Reference D_RG (wild type)** | G_RG | -1.81 | -3.20 to -0.41 | 0.011 |
| | G_KR | 0.58 | -0.57 to 1.72 | 0.32 |

**B**

| | | **Estimate** | **95% CI** | **P value** |
|---|---|---|---|---|
| Days from symptom onset | | 0.61 | 0.39 – 0.84 | 9.9E-08 |
| Spike 614 & Nucleocapsid 203/204 status: **Reference G_RG** | D_RG | 1.81 | 0.41 – 3.20 | 0.011 |
| | G_KR | 2.38 | 1.24 – 3.52 | 4.51E-05 |

**Results from multivariable linear regression models. n=478 individuals sampled in Sheffield, UK. D_RG = D614/R203/G204; G_RG = G614/R203/G204; G_KR = G614/K203/R204.*
